# Conservation of affinity rather than sequence underlies a dynamic evolution of the motif-mediated p53/MDM2 interaction in teleosts

**DOI:** 10.1101/2023.08.24.554616

**Authors:** Filip Mihalič, Dahiana Arcila, Mats E. Pettersson, Pouria Farkhondehkish, Eva Andersson, Leif Andersson, Ricardo Betancur-R, Per Jemth

## Abstract

The transcription factor and cell cycle regulator p53 is marked for degradation by the ubiquitin ligase MDM2. The interaction between these two proteins is mediated by a conserved binding motif in the disordered p53 transactivation domain (p53TAD) and the folded SWIB domain in MDM2. The conserved motif in p53TAD from zebrafish displays a 20-fold weaker interaction with MDM2, compared to the interaction in human and chicken. To investigate this apparent difference, we tracked the molecular evolution of the p53TAD/MDM2 interaction among ray-finned fishes (Actinopterygii), the largest vertebrate clade. Intriguingly, phylogenetic analyses, ancestral sequence reconstructions, and binding experiments showed that different loss-of-affinity changes in the canonical binding motif within p53TAD have occurred repeatedly and convergently in different fish lineages, resulting in relatively low extant affinities (*K*_D_ = 0.5-5 μM). However, for eleven different fish p53TAD/MDM2 interactions, non-conserved regions flanking the canonical motif increased the affinity 4 to 73-fold to be on par with the human interaction. Our findings suggest that compensating changes at conserved and non-conserved positions within the motif, as well as in flanking regions of low conservation, underlie a stabilizing selection of “functional affinity” in the p53TAD/MDM2 interaction. Such interplay complicates bioinformatic prediction of binding and call for experimental validation. Motif-mediated protein-protein interactions involving short binding motifs and folded interaction domains are very common across multicellular life. It is likely that evolution of affinity in motif-mediated interactions often involves an interplay between specific interactions made by conserved motif residues and non-specific interactions by non-conserved disordered regions.

## INTRODUCTION

The interaction between p53 and its regulator the ubiquitin ligase MDM2 is a well-studied system due to the central role of p53 as a master cell cycle transcription factor (1, 2). The interaction between the proteins is mediated by a conserved binding motif in the intrinsically disordered N-terminal transactivation domain (TAD) of p53 and a SWIB domain in MDM2 (**Fig. 1**). The interaction leads to MDM2-catalyzed ubiquitination and proteasomal degradation of p53. Phosphorylation of p53TAD increases affinity for the general transcriptional coactivators CBP/p300 and decreases affinity for MDM2 (3–5). This phosphorylation-dependent activation of p53 leads to transcription of several genes, including the gene encoding MDM2, resulting in a negative feedback loop and fluctuating low levels of p53 under normal conditions. Bioinformatic and experimental analyses suggested that the p53-MDM2 regulation is an ancient mechanism, already present in the common ancestor of all metazoans (6–9). We recently investigated the deep evolution of affinity between p53TAD and MDM2, and found that the interaction between the conserved binding motif in p53TAD and MDM2 SWIB is relatively weak in invertebrates, ranging from not measurable to *K*_D_ = 15 μM in the mollusk *Mytilus trossulus* (10). Furthermore, the affinity was very low (*K*_D_ = 250 μM) in the non-jawed vertebrate arctic lamprey (*Lethenteron camtschaticum*), whereas in other vertebrates such as human and chicken (*Gallus gallus*) the affinity of the interaction is in the 100 nM range. We found that a reconstructed p53TAD from the ancestor of fishes and tetrapods bound with high affinity to its contemporary MDM2 as well as to extant MDM2s from human and zebra fish (*Danio rerio*). However, the extant *D. rerio* interaction between a peptide corresponding to the conserved binding motif in p53TAD*_D.rerio_* (residues 15-26) and MDM2*_D.rerio_* displayed no affinity in our binding assay due to an apparent insertion of an asparagine residue into the binding motif at position 26, which destabilizes the binding interface. The addition of leucine (p53TAD^15-27^), the next residue of the motif, partially restored the canonical hydrophobic triad in the motif (F_19_xxxW_23_xxN_26_L_27_) and resulted in measurable affinity, but still approximately 20-fold weaker than the p53TAD-MDM2 interactions of human, chicken, and the fishes/tetrapods ancestor. Apart from the Asn_26_ insertion, the lower affinity of the *D. rerio* is also due to the absence of Thr_18_, a helix N-cap residue present in most vertebrate p53TADs and whose phosphorylation decreases affinity for MDM2 and increases affinity for CBP/p300 (3, 5, 11).

**Figure 1.**
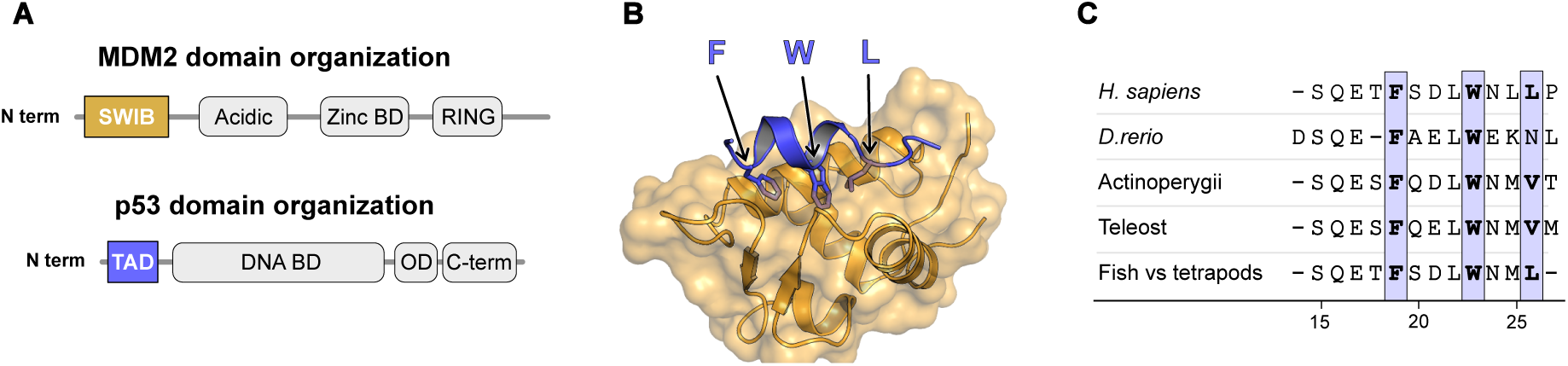
Structure of the human MDM2/p53TAD complex. (A) Schematic domain architecture of MDM2 and p53. (B) Crystal structure of the human complex between the SWIB domain of MDM2 (gold) and a peptide corresponding to the conserved binding motif in p53TAD (blue) (PDBid: 1ycr) (12). Arrows highlight the conserved hydrophobic triad where the residues are shown as sticks. (C) Sequence alignment of reconstructed and extant binding motifs of p53TAD. To facilitate comparison, the site numbering used here follows the human p53TAD binding motif, from Ser15 to Pro27. The previously reconstructed maximum likelihood sequence for the most recent common ancestor of bony fishes and tetrapods (10) is indicated in the alignment as “Fish vs tetrapods.”

In the present study, we further investigate this intriguing apparent loss of affinity in *D. rerio* p53TAD by performing a comprehensive phylogenetic analysis of p53TAD for Actinopterygii (ray-finned fishes), combined with experiments to determine the affinity between p53TAD and the SWIB domain of MDM2 from representative extant fish families, and for reconstructed p53TAD and MDM2 variants from ancestral fishes. A fascinating picture emerges, in which loss-of-affinity mutations in the conserved MDM2-binding motif of p53TAD have occurred several times in different fish lineages. We also find that the loss of affinity in p53TAD is compensated by the C-terminal flanking region of the conserved motif. Our analyses show how regions outside of the conserved binding motif promote binding and contribute to an extraordinarily dynamic evolution of the p53-MDM2 interaction. These findings demonstrate a high plasticity of motif-mediated interactions involving transactivation domains such as p53TAD and the uncertainty of bioinformatic inference of binding without experimental validation. It is likely that similar mechanisms of affinity modulation by flanking regions are widespread in motif mediated interactions.

## RESULTS

### Phylogenetic analysis of fish p53TAD

To investigate the evolution of p53TAD in ray-finned fishes, we collected p53 sequences from 51 species of Actinopterygii. We also included three other vertebrates as outgroups in the sequence alignment: human (*Homo sapiens*), coelacanth *Latimeria chalumnae* and Australian ghostshark (*Callorhinchus milii*). In several species, we found two distinct p53 proteins, and phylogenetic analyses suggested that they are paralogs originating in the third whole genome duplication in teleosts (13), the dominant clade within Actinopterygii. In salmonids, that underwent a fourth genome duplication (14), we found three paralogs of p53. In cases where the phylogeny of the two p53 paralogs was uncertain from sequence analysis alone, such as *Anguilla anguilla* (European eel), which diverged early from other teleost fishes, the identity of the paralog was confirmed by its local chromosomal context, assessed from the genome databases at ENSEMBL and NCBI. Paralogs assigned to “Class II” were flanked by *gps2* in the sense direction, on the upstream side and an anti-sense sequence of an E3 ubiquitin-protein ligase followed by a *capga* homolog on the downstream side. “Class I” paralogs had variable downstream contexts, but were always flanked by *slc2a4* in anti-sense direction on the upstream side. The sequence variation within p53TAD is too large and with too many insertions and deletions (indels), which makes accurate estimation of homology and a reliable gene tree inference challenging. Instead, we employed a reference species phylogeny (15, 16) as the analytical framework. This approach not only mitigated these challenges but also facilitated the identification of ancestral changes in the amino acid sequence across different branches of the fish tree of life.

Amino acid sequences were first aligned using Clustal Omega (17), followed by manual alignment curation. Indeed, a high-confidence alignment of the entire p53TAD is difficult to obtain due to the extensive changes in most parts of the sequence, including both point mutations and indels. Nevertheless, the obvious limitations of the alignment also demonstrate the very dynamic evolution of p53TAD that has occurred along different lineages (**Fig. 2**). The p53TAD alignment was used together with the established fish phylogeny to reconstruct the conserved binding motif in p53TAD at different ancestral nodes using maximum likelihood in MEGAX (**Supplementary Spreadsheet 1, Fig S1**). Note that we show a reconstruction of the entire p53TAD in supplemental information, but it is only the 12-residue canonical binding region, which interacts with MDM2, as well as the N-terminal region of the DNA-binding domain that served as a reference point, that can be reconstructed with any confidence. (**Supplementary Text File 1** presents a list of posterior probabilities for each position in each node). In addition, a second putative transactivation domain motif is conserved among most teleosts (**Fig. 2**). The reconstructed sequence of this second motif for the ancestral teleost is FDEKLFE (**Supplementary Spreadsheet 1**) and the consensus motif for the teleost sequences in Figure 2 is FDENLFE (determined with Jalview (18)). A reconstruction of ancestral maximum likelihood sequences was also performed for the folded SWIB domain of MDM2 (**Supplementary** Fig. 2**, Supplementary Spreadsheet 2, Supplementary Text File 2**). To account for uncertainty in the reconstruction we performed experiments with alternative reconstructed variants (AltAll) where residues with a probability below 0.9 were replaced by the second most likely residue (19) (**Supplementary Spreadsheet 3 and 4**). Based on the available sequence data and the reference phylogeny, there are several crucial historical events of molecular evolution that can be established with high confidence.

**Figure 2.**
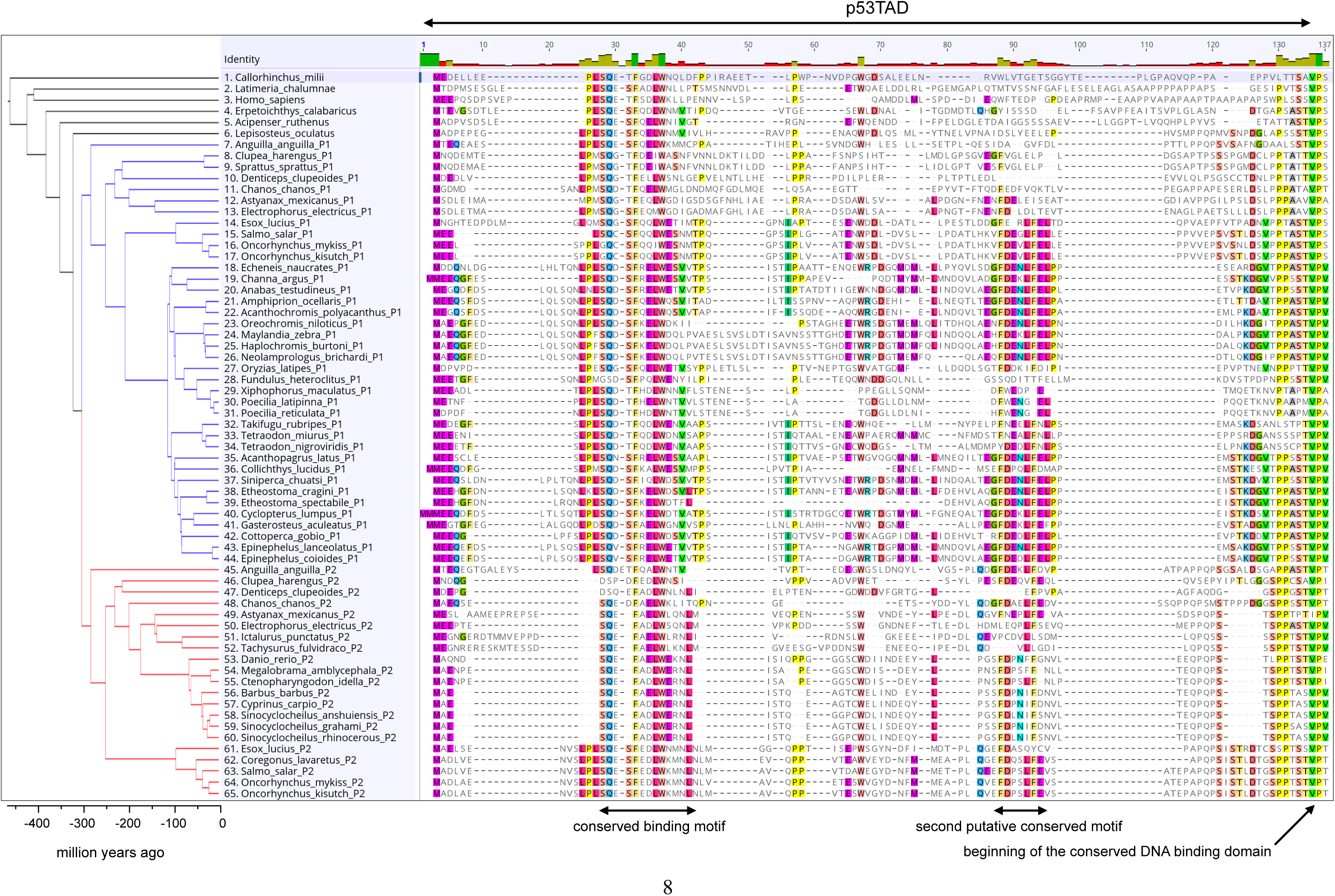
Sequence alignment of p53TAD mapped onto the fish phylogeny. We included 51 extant species representing some of the major lineages of fishes and for which high-quality sequencing data were available. The blue and red branches represent the split between the two paralogs p53TADP1 and p53TADP2, respectively, in teleosts (node 125). After the split, p53TADP1 and p53TADP2 have evolved in parallel in each lineage. Node numbers refer to the ancestral reconstruction analysis (**Supplementary Spreadsheet File 1**).

### Evolution of the binding affinity between p53TAD canonical motif and MDM2 in fishes

To trace the evolution of affinity of the p53TAD binding motif across Actinopterygii species we expressed and purified reconstructed and extant MDM2 SWIB domains and measured the affinity using a fluorescence polarization-based method. We included four ancient (ancestral Actiopterygenii, teleost, paralog 1 and paralog 2, respectively), 12 extant MDM2s and p53TAD peptides containing the conserved binding motif (corresponding to human p53 residues 15-27, p53TAD^15-27^, **Fig. 1**). The extant sequences were from eight fishes representing different evolutionary lineages in the phylogenetic tree, and four paralogs from three of these species. Additionally, we included the human MDM2/p53TAD interaction for reference (**Table 1**, **Fig. 3, Table S5**).

**Figure 3.**
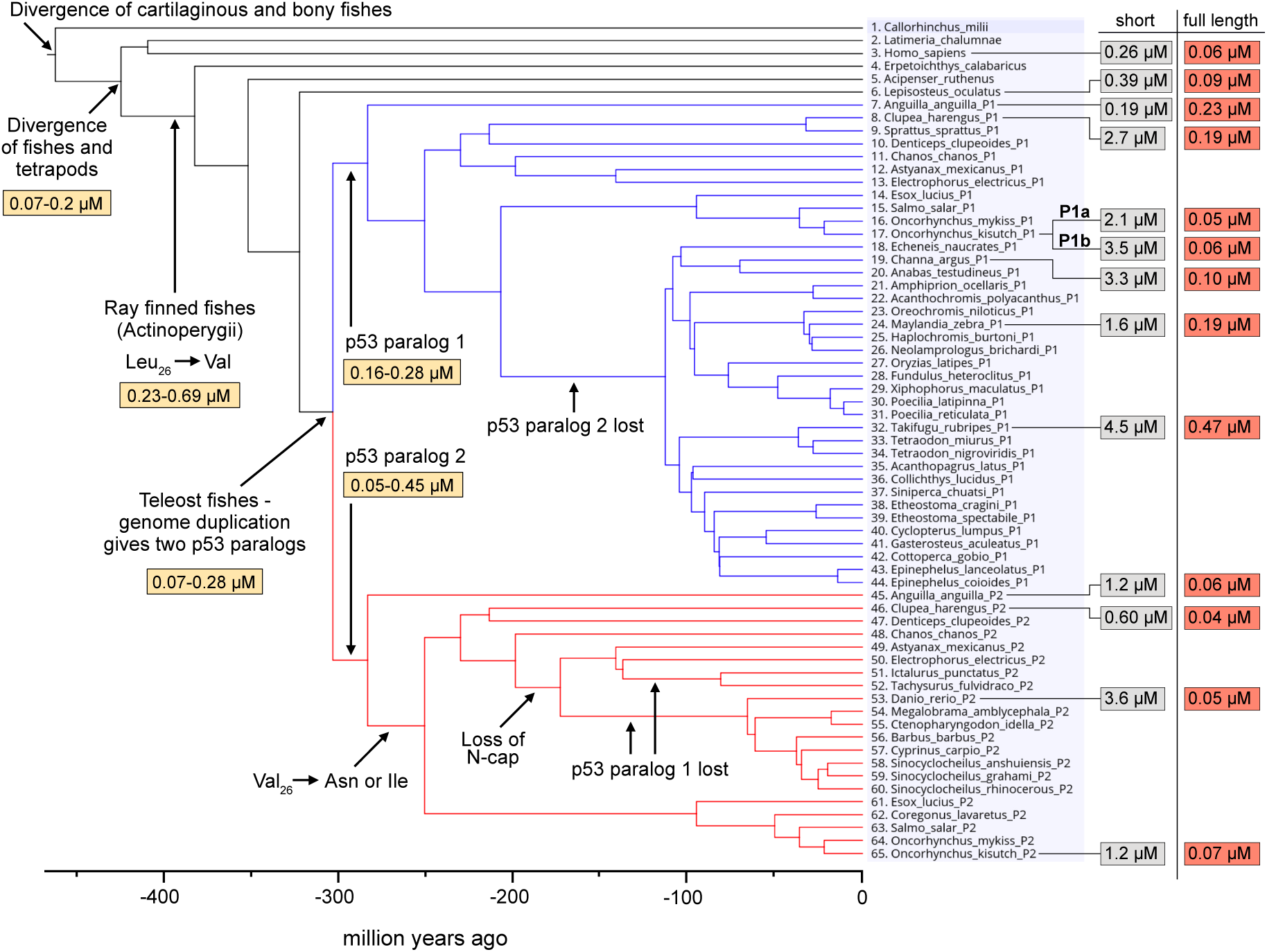
Affinities of p53TAD/MDM2 interactions. Affinities (*K*D values) of extant and ancient p53TAD^15-27^/MDM2 for different fishes mapped onto the phylogenetic tree. Key events that are discussed in the text are shown. Affinities for the reconstructed p53TAD^15-27^/MDM2 are highlighted in yellow, the affinities of p53TAD^15-27^/MDM2 for present day fish species in gray and for the p53TAD^full^ ^length^/MDM2 in red. See **Table 1** and **Supplementary Table 1** for errors.

**Table 1.**
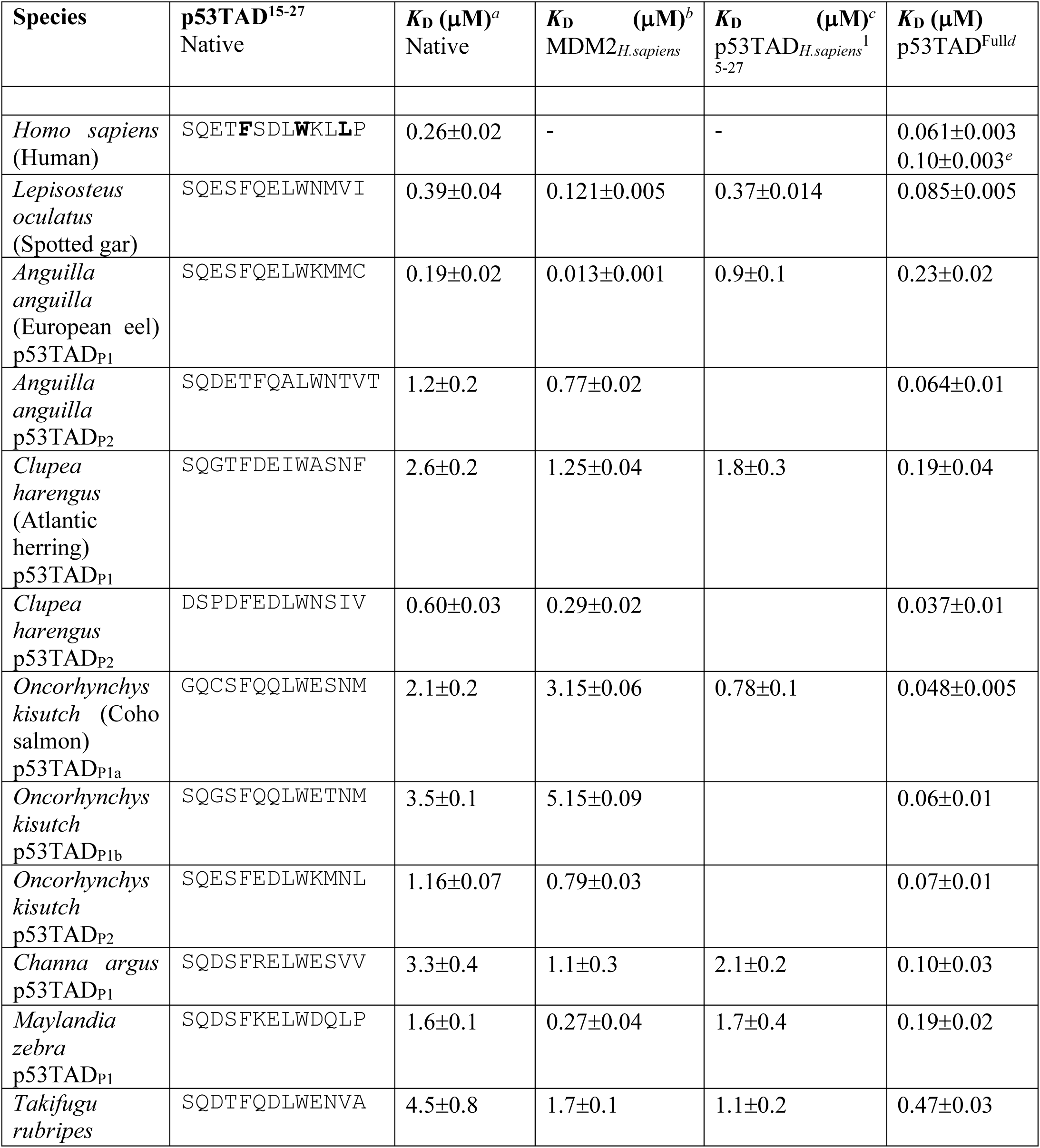
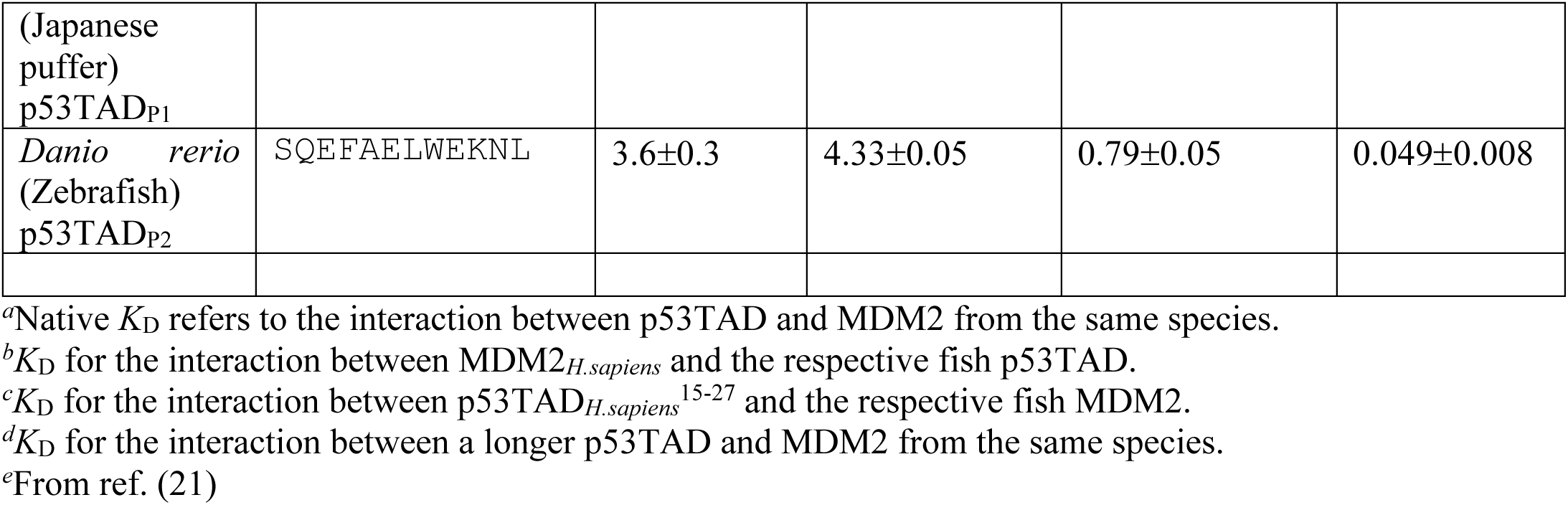
Affinities for extant MDM2/p53TAD interactions. *K*_D_ values were measured using a fluorescence polarization-based displacement assay. The complex between MDM2 and FITC- labeled p53TAD_Human_^15-27^ was displaced by unlabeled p53TAD^15-27^ or full-length p53TAD^Full^ from different species and the IC_50_ value was used to calculate *K*_D_.

Previous reconstruction of the p53TAD canonical binding motif from the common ancestor of fishes/tetrapods (10) suggested the presence of all three hydrophobic residues (in bold) that point into the binding pocket and make interactions with MDM2 in human p53TAD: S_15_Q_16_E_17_T_18_**F_19_**S_20_D_21_L_22_**W_23_**N_24_M_25_**L_26_** (**Fig. 1**). While the Phe_19_ and Trp_23_ are strictly conserved among Actinopterygii, there is considerable variation with regard to the third hydrophobic residue (Leu_26_) and the following position 27, which was not included in the previous reconstruction of p53TAD_Fishes/Tetrapods_ (10). Here, we included both residues in the experiments since residue 27 contributes to affinity in p53TAD peptides derived from extant *D. rerio* (10). In the p53TAD_Actinopterygii_^ML^ motif, the ancestral Leu_26_ in p53TAD_Fishes/Tetrapods_^ML^ had been replaced by Val_26_. This, along with two other substitutions (**Fig. 1C**) resulted in an affinity of 690 nM towards reconstructed maximum likelihood MDM2_Actinopterygii_^ML^ (**Fig. 3**, **Supplementary Table 1**). This affinity is lower than the affinity between the ancestral fishes/tetrapods proteins (*K*_D_ = 70-200 nM). Indeed, Leu_26_→Val is a loss-of-affinity mutation, as shown by the 5 to 10-fold increase in affinity of p53TAD_Actinopterygii_^ML^ and p53TAD_Teleost_^ML^ peptides with the reverse Val_26_→Leu mutation (**Supplementary Table 1**). The reconstructed AltAll sequences for the ancestral Actinopterygii complex display slightly higher affinities than the maximum likelihood variants, yielding a *K*_D_ = 230 nM for the MDM2_Actinopterygii_^AltAll^ and p53TAD_Actinopterygii_^AltAll^ interaction.

Val_26_ is conserved in p53TADs of extant non-teleost Actinopterygii and in paralog 1 of most percomorphs (18-44 in the gene tree, **Fig. 2**). The reconstructed p53TAD at node 125, representing an ancestral teleost p53TAD before the genome duplication, has a posterior probability of 0.60 for Val_26_ and 0.39 for Ile_26_ (**Supplementary Spreadsheet 3, Fig. S1**). Despite this uncertainty, a comparison of affinities (**Fig. 3, Supplementary Table 1**) of the peptides representing the ancestral p53TAD_Teleost_, p53TAD_P1_ and p53TAD_P2_ (see below) suggests that higher affinity possibly evolved in the ancestral teleost as compared to the ancestral ray-finned fish by a change from Val_26_Thr(Ala)_27_ to Val(Ile)_26_Met(Leu/Val)_27_, with AltAll amino acids in parenthesis. However, if the alanine was present at position 27 in p53TAD_Actinopterygii_, the ancestral Actinopterygii and teleost complexes were of similar affinity.

The teleost-specific whole genome duplication created two paralogous p53 genes with separate evolutionary histories (**Fig. 2, Fig. S1**). The reconstructed last common ancestor of paralog 1 (p53TAD_P1_, node 104) and paralog 2 (p53TAD_P2_, node 124) are both very similar to p53TAD_Teleost_, and the three reconstructed binding motifs have the same ambiguity with regard to Val_26_/Ile_26_. However, since Val_26_ is spread over the p53TAD_P1_ clade defined by node 101, and is also present in p53TAD_P2_ of *A. anguilla*, the most likely scenario is that Val_26_ is ancestral for p53TAD_P1_, p53TAD_P2_ and p53TAD_Teleost_ and that Ile26 is derived later in certain lineages. Assuming that Val_26_ is ancestral, this residue is largely conserved in most percomorphs (species 18-44) in p53TAD_P1_. Position 27 is very uncertain for p53TAD_Teleost_, with around 25% probability for either Met, Leu or Val, reflecting the different residues at this position in extant fishes. Thus, the ancestral Val_26_ and the uncertain residue at position 27 have been replaced by different residues in non-percomorph teleosts, a paraphyletic group encompassing extant species 8-17, including *Clupea harengus* (herring) (Asn_26_Phe_27_), *Denticeps clupeoides* (denticle herring) (Leu_26_Gly_27_), *Electrophorus electricus* (electric eel) (Ile_26_Gly_27_), and the salmonids (Asn_26_Met_27_) (**Fig. S3**). In p53TAD_P2_ we observe further intriguing modifications at the end of the binding motif. Either in ancestral node 123, after the divergence of the p53TAD_P2_ branch leading to *A. anguilla*, or, alternatively, at least three times in separate branches (**Fig. S4**) there is a change of Val_26_→Asn. This mutation to Asn_26_ is particularly notable since it has occurred in certain p53TAD_P1_ lineages as well (salmonides and *Sprattus sprattus*), thus representing a recurrent loss-of-affinity change. Extant *A. anguilla* p53TAD_P2_ retains Val_26_ either as a result of back mutation or conservation. *Clupea harengus* (Ile_26_Val_27_) and *Chanos chanos* p53TAD_P2_ (Ile_26_Thr_27_) have gained subsequent mutations at the end of the motif. We note here that it is difficult to judge whether the described changes at positions 26-27 are due to point mutation or indels, as several residues in the region are different among extant fishes, obscuring the historical order of events (**Fig. S3, Fig. S4**).

To further rationalize determinants of affinity we aligned the p53TAD binding motifs and ranked them based on *K*_D_ values for native MDM2 (**Fig. S5a**) and for MDM2_H.sapiens_ (**Fig. S5b**) which clearly showed that non-conserved residues such as those at position 24 and in particular 25 can affect affinity. To some extent, in the case that the residue in position 26 is not hydrophobic, position 27 will also influence binding. Interestingly, Met_25_ is recurrent in high-affinity p53TAD peptides, in particular with Leu_26_, Ile_26_ or Met_26_ in the adjacent position, but also with Val_26_. Met_25_ was likely present in the ancestral p53TAD_Actinopterygii_ and p53TAD_Teleost_. Its subsequent mutation in node 103, *i.e.*, all p53TAD_P1_ except *A. anguilla*, and in most p53TAD_P2_ (except the salmonides) is another example of a recurrent loss-of-affinity change. The N-cap at position 18 stabilizes the helix formed by the conserved 12-residue motif upon binding to MDM2, via a hydrogen bond to Asp_21_ in the human complex (12). The ancestral p53TAD_Teleost_ Ser_18_ was likely replaced in node 118 (p53TAD_P2_) by Asp_18_, which could potentially also function as N-cap (**Fig. 2**). However, Asp_18_ cannot act as a phosphorylation switch regulating MDM2 and CBP/p300 affinity and was subsequently deleted in the common ancestor of otophysans (p53TAD_P2_, species 49-60, including *D. rerio*). Experiments using different p53TAD*_D.rerio_* peptides previously demonstrated that this is another loss-of-affinity mutation (10).

We also observed complete gene loss in some taxa. In fact, p53_P1_ was apparently lost twice, once in an ancestor of cypriniforms (species 53-60, including *D. rerio*) and once in the lineage leading to non-cypriniform otophysans (species 51-52). The gene encoding p53_P2_ was lost in the common ancestor of the percomorph lineages Ovalentaria, Carangaria and Anabantaria (species 18-31; node 88). Thus, we found both paralogs, p53_P1_ and p53_P2_, in a limited number of species including *A. anguilla*, *C. harengus* and the salmonid family. Clearly, an alternative explanation is that sequence data are incomplete and that we miss some paralogs. However, the clustering of sequences (or lack of sequences) in the phylogenetic tree supports a scenario where the paralogs were indeed lost, as it is unlikely that potential assembly errors would be systematic in such fashion. Furthermore, we investigated chromosome locations surrounding the characteristic flanking genes but were unable to detect additional p53 homologs.

Intriguingly, the affinities of the extant complexes generally had relatively weak affinities across the fish tree between native p53TAD^15-27^ binding motifs and MDM2 (*K*_D_ = 0.2-5 μM), as compared to the ancestral interactions (0.05-0.7 μM) and the human interaction (*K*_D_ = 0.26 μM) (**Fig. 3**). For a direct comparison of the different fish p53TADs and MDM2s, we also measured (*i*) the affinity between p53TAD_Human_^15-27^ and the fish MDM2s, and (*ii*) the affinity between fish p53TADs and human MDM2. The trend was the same with relatively low affinities for fish p53TADs as compared to the human interaction. Moreover, the affinities were in general slightly higher when either human p53TAD or human MDM2 was in the complex as compared to the native fish interaction (**Table 1**).

Our experimental results and the sequence alignment of fish p53TADs suggest an intriguing scenario: the affinity between the canonical binding motif in p53TAD and MDM2 appears to first have decreased in the ancestral Actinopterygii p53TAD (Leu_26_→Val), then increased by changes at position 27 to a large hydrophobic residue in the ancestral p53TAD_Teleost_ and then – for example, via deletion of Asp_18_, changes at Met_25_, or Val_26_→Asn – decreased again such that ten of the twelve investigated present-day canonical motifs display *K*_D_ values in the range of 0.6-4.5 μM across teleost fishes.

### Flanking regions increase the affinity between p53TAD and MDM2

Despite the relatively low affinity, MDM2 is crucial for regulating p53 levels in *D. rerio* as demonstrated by knockout experiments (20). Therefore, we expressed, purified, and subjected a longer p53TAD construct from *D. rerio* to binding experiments (**Fig. 4A**). Surprisingly, we found that this construct, p53TAD*_D.rerio_*^10-75^, displayed more then 70-fold higher affinity (*K*_D_ = 49 nM) than the p53TAD*_D.rerio_*^15-27^ and even higher affinity than human and chicken p53TAD/MDM2 interactions. For the human complex, a corresponding p53TAD_Human_^13-61^ increased the affinity only 1 to 4-fold (depending on ionic strength) compared to the shorter p53TAD_Human_^15-29^, as determined by kinetic experiments (21) and confirmed in this study (4-fold by fluorescence polarization, **Fig. 3**, **Supplementary Spreadsheet 6**).

**Figure 4.**
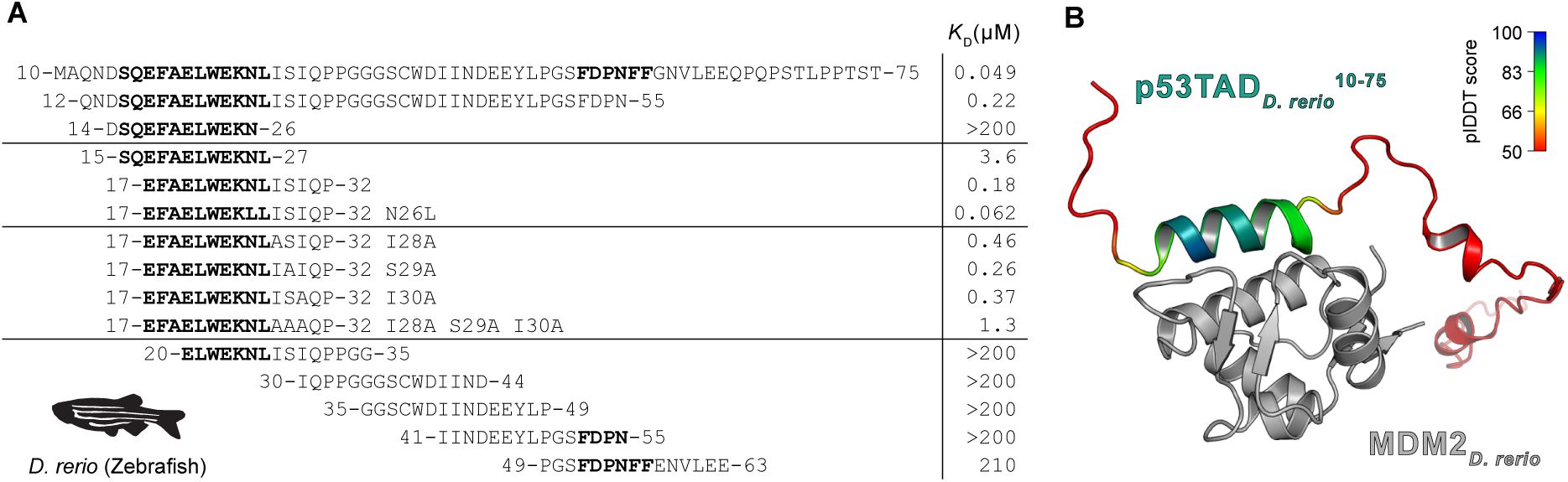
Affinities between p53TAD peptides and MDM2 from *Danio rerio*. (A) The overlapping peptides tiling a 51-residue stretch of p53TAD*D.rerio*. To facilitate comparison, the numbering starts from the conserved motif and the residues of human p53TAD, F19xxxW23xxL26. *K*D values for the peptide fragments were determined by the fluorescence polarization assay. Distal residues as well as Ile28Ser29Ile30 contribute to the affinity for MDM2 (B) Colabfold prediction of the interaction between *D. rerio* MDM2 and p53TAD^10-75^. The MDM2 is in gray, and the p53TAD^10-75^ is colored according to the pIDDT, where blue indicates highest and red lowest prediction confidence (pIDDT<50).

To pinpoint the residue(s) contributing to the higher affinity, six overlapping peptides corresponding to p53TAD*_D.rerio_*^12-63^ were used in fluorescence polarization competition experiments. The peptide p53TAD*_D.rerio_*^17-32^ containing the conserved binding motif and the additional residues ISIQP fully recapitulated the affinity of p53TAD*_D.rerio_*^12-55^ (*K*_D_ = 180 nM, **Fig. 4A**), but not of the full-length p53TAD*_D.rerio_*^10-75^. The remaining shorter peptides displayed negligible affinity. We additionally observed that the Uniprot protein sequence database contains two different *D. rerio* p53 sequences under entries G1K2L5 and P79734, the latter marked as canonical sequence and lacking the Ser_29_. This alternative p53TAD*_D.rerio_*^17-32^ (P79734), ending with IIQPP, bound slightly weaker (*K*_D_ = 580 nM) than ISIQP. We then replaced each of the residues Ile_28_Ser_29_Ile_30_ with Ala. The combined effect of each point mutation (7.5-fold reduced affinity) was identical within experimental error to the triple Ala mutant (AAAQP) (7.7-fold), showing that several residues close to the motif contribute additively to the increase in affinity (**Fig. 4A**).

To rationalize this result we performed Colabfold (22) predictions of the *D. rerio* MDM2/p53TAD^10-75^ complex and observed that the algorithm predicted an extension of the p53TAD binding helix including residues ISIQP (Fig. 4B). Such an extended helix is not predicted for the human complex (**Fig. S6**). Importantly, in p53TAD*_H.sapiens_* the canonical motif is immediately followed by a helix-breaking proline (position 27). Previous experiments on the human complex involving a Pro_27_→Ser substitution showed that the helicity increased and the affinity decreased 10-100 fold (23), an effect we confirmed here with a Pro_27_→Ala variant of p53TAD*_H.sapiens_*^17-32^ (**Fig. S7**, **Supplementary Spreadsheet 6**).

Inspired by the data for the *D. rerio* complex, we attempted the expression and purification of longer p53TADs from the same extant fishes where we already measured the affinity for the shorter p53TAD^15-27^ constructs, to assess whether the high affinity of the longer p53TAD is a universal phenomenon across the fish clade. These longer p53TAD^Full^ ^length^ constructs consisted of the entire region from the first amino acid of p53 to the beginning of the conserved DNA-binding domain, which is the C-terminal part of the alignment (**Fig. 2**, **Supplementary Spreadsheet 5**). We were able to express and purify full length p53TADs for all twelve fish interactions. Remarkably, in eleven cases the affinity increased considerably (4 to 73-fold, **Table 1**, **Fig 3**, **Fig. 5**) resulting in *K*_D_ values in the same range as that of the human complex (∼100 nM). Inspection of the flanking C-terminal sequences shows a low degree of conservation suggesting that different combinations of amino acid residues can promote increased affinity (**Fig. S8**). Additionally, the Colabfold predictions identified extended helices for several but not for all cases of bound p53TADs from different fishes suggesting alternative mechanisms for increasing the affinity, as compared to *D. rerio* p53TAD/MDM2 (**Fig S9**).

**Figure 5.**
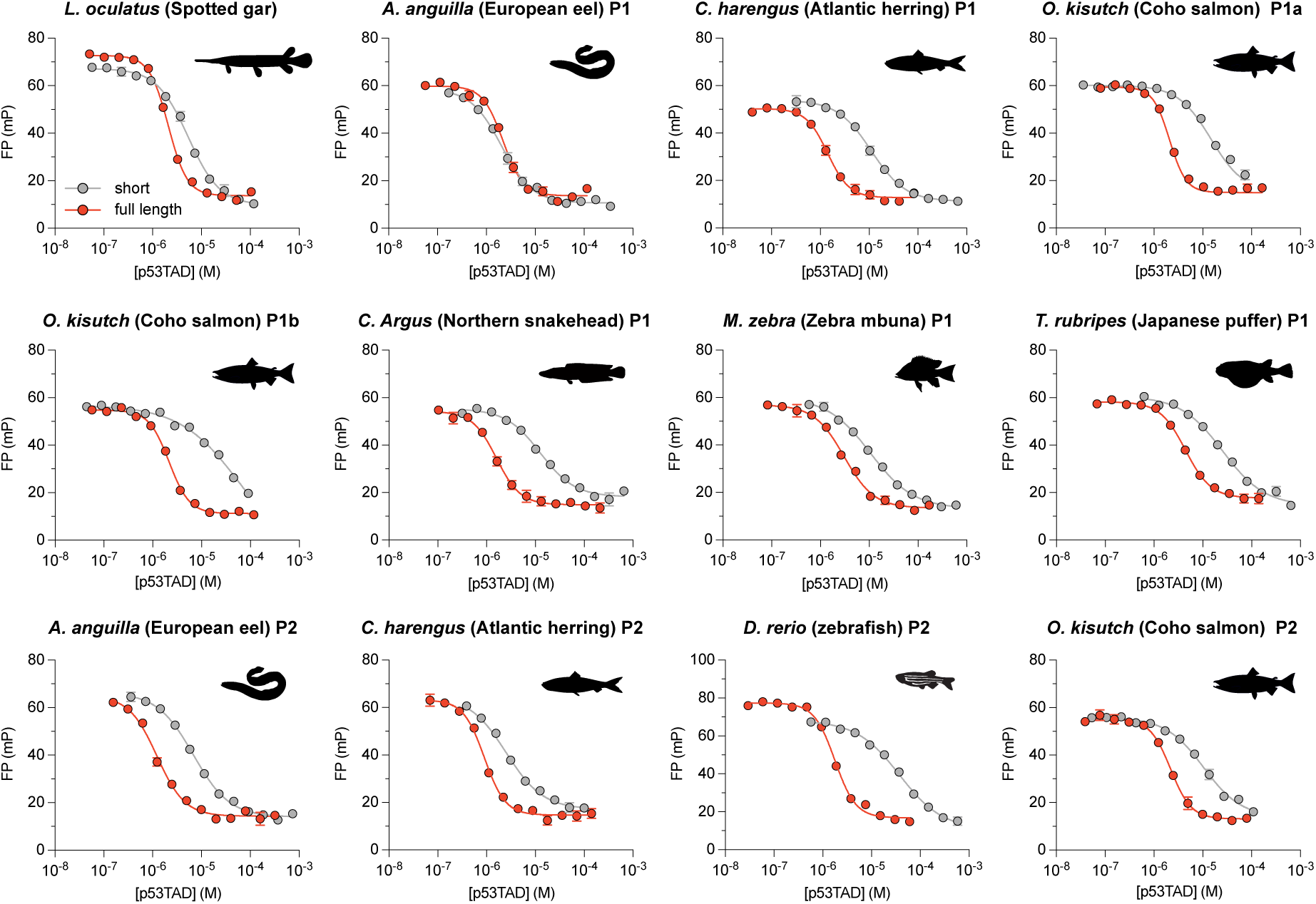
Affinity of short and full length p53TAD of different extant fishes measured by fluorescence polarization. Affinity is higher for full length p53TAD constructs (Fig. 2, **Supplementary Spreadsheet 5**) as compared to the short p53TAD^15-27^, often by an order of magnitude, with the exception of *A. anguilla* p53TADP1. FP, fluorescence polarization in millipolarization units (mP).

## Discussion

The gene encoding an ancestral p53-like protein was duplicated twice in an early vertebrate around 450 million years ago (Mya), before the divergence of cartilaginous fishes, bony fishes, and tetrapods, in two whole genome duplications (24–27). From these events, three paralogs remain in extant vertebrates, namely p53, p63 and p73. Vertebrate p53 evolved to become a key cell cycle regulator, keeping its ancestral regulation by the ubiquitin ligase MDM2 (1, 2). The evolution of the molecular interaction between a conserved motif in the transactivation domain of p53-family proteins and MDM2 has been very dynamic with reduced affinity in some animal lineages (10) and even complete loss of the interaction domains (28). We previously reconstructed p53TAD and MDM2 from the last common ancestor of fishes and tetrapods (Fig. 1). In the present study we follow the branches down the Actinopterygii lineages to investigate the evolution of affinity between p53TAD and MDM2 in fishes. Interestingly, loss-of-affinity mutations in the binding motif have been selected for on several occasions in different fish lineages: Leu_26_→Val (400 Mya), Met_25_→Ser (250 Mya, p53TAD_P1_), Val_26_→Asn (250 Mya or later, p53TAD_P2_), deletion of an N-cap residue (180-200 Mya, p53TAD_P2_), and Ile_26_→Asn (40-90 Mya, p53TAD_P1_) (**Fig. 2, Fig. S3, Fig. S4**). In addition, complete loss of the paralogs occurred: p53_P1_ (70-180 Mya, likely in two separate branches), and p53_P2_ (120-210 Mya).

In a larger context, a wealth of data has demonstrated the importance of short binding motifs for protein-protein interactions (29). The interaction between a typical binding motif and an interaction domain usually involves a “short linear interaction motif” of only 3-10 amino acid residues, which provides a functional affinity with *K*_D_ values in the nM-μM range. However, recent data suggested that flanking disordered regions of the binding motif may modulate affinity (30–32). Activation domains are enriched in hydrophobic and negatively charged residues, where the latter likely contribute to exposure of hydrophobic motifs (33). Whilst certain residues (aromatic, Leu) are vital, it has been proposed that the composition, rather than exact sequence, plays a more important role for transcriptional activity (33–35). However, our data show that details in the motif could indeed matter. For example, it would be hard to predict that *A. anguilla* p53TAD_P1_ is the sole high-affinity extant teleost motif among those included in the study, with a positively charged Lys next to Trp_23_ and a Met at position 26.

Our present work on the p53/MDM2 interaction in ray-finned fishes provides a striking example of the potential role of rapidly evolving flanking regions, where the C-terminal flanking regions from eleven out of twelve different Actinopterygii p53TADs increase the affinity 4 to 73-fold as compared to the canonical conserved binding motif. In the human p53TAD/MDM2 complex, with a relatively high affinity for the canonical motif, only a 4-fold increase was observed. Indeed, also for the fish complexes, the lowest influence from flanking regions was found in the two cases with highest affinity of the motif (**Fig. 3**). It is important to note that apart from MDM2, several other proteins interact with p53TAD (2). Thus, the dynamic evolution of the p53TAD sequence, and its interaction with MDM2, is likely influenced by other proteins and interactions that may fine tune the regulation of p53.

Clearly, residues 25-27 influence affinity, and this region samples different combinations of residues over evolutionary time, possibly contributing to adaptation when a new niche is explored. These and other loss or gain-of-affinity mutations could be compensated by additional changes in the C-terminal flanking region to maintain affinity. Since reconstruction of the C-terminal part of p53TAD is not possible, we can only speculate about the order of events. One possible scenario is that the extensive mutations occurring in the C-terminal part first led to increased affinity between the proteins, resulting in a suboptimal but manageable high affinity. Compensatory mutations in the binding motif could then restore a lower affinity. It is harder to envision initial loss-of-affinity mutations in the binding motif since a 10-fold decrease in affinity would likely lead to severe effects on p53 regulation, impaired fitness and elimination of the variant by negative selection. Initial mutations leading to changes in concentration of proteins could also be the cause of subsequent mutations in the binding motif of p53TAD to restore fitness. In this context it is interesting to consider Pro27, which is present in extant mammals, birds and frogs (10) and therefore likely present in the ancestral tetrapod. This residue lowers the affinity of the human interaction as shown by the Pro_27_→Ser (23) or Pro_27_→Ala mutation (**Fig. S7**), and contributes to a functional affinity in the range of 100 nM for the human and chicken interactions. Thus, we speculate that the affinity of an ancestral fish/tetrapod complex became too high and was attenuated in the tetrapod lineage by selection of a helix-interfering proline, while in fishes, different changes at position 26 and 27, and also further C-terminally, modulated affinity. Importantly, the Pro_27_→Ala mutation illustrates how easily the affinity could increase in the human interaction and therefore, by inference, that there is a stabilizing selection for functional affinity rather than for sequence conservation, involving purifying selection against higher affinity. Similar conclusions were reached from experiments in yeast involving poorly conserved disordered regions in MAP kinase signaling (34).

As a general mechanism, flanking regions can buffer against changes in the binding motif and *vice versa*, as recently noted for yeast general regulatory factors (36). Our data suggest that many of the relatively weak interactions observed between binding motifs and interaction domains in general (*K*_D_ in the high μM-low mM range) may be stronger due to non-specific interactions provided by flanking regions. A picture is emerging where context in terms of flanking regions and even supertertiary structure (37) prove important for a functional affinity in motif-mediated interactions between short disordered interaction motifs and globular domains (30, 31, 38).

## Materials and methods

### Reconstruction of ancestral sequences

We aligned 62 p53TAD sequences, including both paralogs p53TAD_P1_ and p53TAD_P2_, from 51 Actinopterygii species initially using ClustalO (17), followed by manual curation. In uncertain cases the identity of the respective paralog was confirmed by chromosomal location. This was done by manual inspection of the ENSEMBL (release 104) and NCBI (accessed Nov 2021) genome browsers. p53TADs from human, *Latimeria chalumnae* and *Callorhinchus milli* were used as outgroups. In addition to p53TAD, we also aligned 48 MDM2 SWIB domain sequences. Ancestral sequence estimation for both p53TAD and MDM2 SWIB was conducted in MEGAX v10.1.8 (39) using the JTT+G+I model of sequence evolution (with 5 gamma categories) and the maximum likelihood reconstruction method (**Supplementary Spreadsheets 1 and 2**). The reconstructions used both the sequence alignments and the species phylogeny as input (**Fig. 2**). We used the time-calibrated phylogenomic tree estimated by Hughes et al. (16), after pruning out all non-target species in their tree and manually grafting a few missing species for which we could obtain p53TAD or MDM2 SWIB domain sequences. Missing species grafting followed the phylogenetic placement and ages of these taxa in the fish tree inferred by Rabosky et al. (40). To accurately reconstruct ancestral sequences for the different p53TAD paralogs examined, the teleost whole genome duplication event was accounted for by manually duplicating the teleost clade in the phylogenomic tree at 303 Mya, which is within the timeframe estimated for the duplication event by molecular phylogenetic studies (41).

While the species were overall similar in the p53TAD and MDM2 alignments, there were differences in cases where a reliable MDM2 sequence could not be found. For example, we used *Coregonus lavaretus* p53TAD_P2_ and *C. clupeaformis* MDM2 SWIB domain. In other cases, we did not replace the MDM2, thus ending up with fewer species in total. However, extant fish families were well represented in both reconstructions. For MDM2, the N-terminus of reconstructed node 94 was edited due to too few sequences at positions 4 and 5 (gaps were inserted). Due to the uncertainty and for consistentcy purposes with a previous study (10), we used expression constructs starting at SQ. In addition, the end of the sequence for reconstructed node 94 is also very uncertain because of few fish sequences in the alignment. We used low-probability AltAll versions of p53TAD and MDM2_Teleost_ to check how robust our experimental results were to errors in the reconstructed sequence, an approach previously described in detail (19). For these AltAll variants, we used a cutoff for the posterior probability for the maximum likelihood residue of < 0.90 and a probability of the second most likely residue of at least 0.10. All residues that fulfilled these criteria were then included in the AltAll variant (**Supplementary Spreadsheet 3** and **4, Supplementary Text File 1 and 2**). If the affinity of the ML and AltAll variants are similar, it is likely that the true ancestral variant displayed a similar affinity.

### MDM2 SWIB and full length p53TAD protein expression and purification

The cDNA encoding a p53TAD from *Danio rerio* with both potential TADs (denoted “full length"; (QNDSQEFAELWEKNLISIQPPGGGSCWDIINDEEYLPGSFDPN) residues 12-55 using the numbering for human p53TAD) was ordered in a prSET vector. This resulted in a C-terminally His_6_-Lipo-Thrombin (cleavage site) tagged sequence. The Human MDM2 and the *D. rerio* MDM2 were ordered in a pSY10 plasmid which resulted in constructs that were N-terminally NusA-TEV-His_6_-PreScission (cleavage site) tagged. All other MDM2 and full length p53 proteins used in this study were ordered in pETM33 vector which upon expression produced N-terminally His_6_-GST-PreScission tagged constructs. The addition of hexa-histidine, Lipo, NusA or GST tags was used to facilitate higher expression levels and enable simplified protein purification. All sequences were ordered from GenScript (Netherlands) and can be found in **Supplementary Spreadsheet 5**.

All constructs were purified as described previously (10). In short, *Escherichia coli* BL21 (DE3) cells (Invitrogen) were transformed with plasmid constructs and grown in 2xYT medium at 37°C until the OD_600_ reached 0.6-0-8. At this point protein overexpression was induced with the additio18°C. In the case of p53TAD*_D.rerio_*^Full^, MDM2*_H.sapiens_* and MDM2*_D.rerio_* the cells were palleted and resuspended in binding buffer (400 mM sodium chloride, 50 mM sodium phosphate, pH 7.8, 10% glycerol). Cell lysis was achieved by sonication and the lysate was clarified by centrifugation at 4°C. Supernatant was filtered and loaded onto a Nickel Sepharose Fast Flow column (GE Healthcare). Column was washed with binding buffer and the proteins of interest were eluted with the binding buffer supplemented with 250 mM imidazole and applied to size exclusion chromatography column Hi load 16/60 Sephacryl S-100 column (GE Healthcare). Fractions of interest were pooled, and the cleavage of tag was induced by the addition of Thrombin or PreScisson proteases over night at 4°C. Final run of size exclusion was employed to remove the tag and produce pure protein.

In the case of other constructs cloned into pETM33 vector, protein expression was done in the same manner. Upon sedimentation of bacteria, they were resuspended in binding buffer 2 (50 mM Tris/HCl pH 7.8, 300 mM NaCl, 10 µg/mL DNase I and RNase, 4 mM MgCl_2_, 2 mM CaCl_2_ and cOmplete EDTA-free Protease Inhibitor Cocktail) and sonicated, followed by centrifugation. Supernatant was filtered, applied to Pierce™ Glutathione Agarose beads (Thermo Scientific), washed with wash buffer (50 mM Tris, 300 mM NaCl, pH 7.8) an eluted with wash buffer supplemented with 10 mM reduced glutathione. Eluted fractions were cleaved over night with PreScission protease to remove the GST tag and the tag was removed by applying of the cleavage reaction onto Nickel Sepharose Fast Flow column. Flowthrough was collected and the final polishing step was performed using a Hi load 16/60 Sephacryl S-100 column. Sample purity after final step of purification was assessed by SDS-PAGE and MALDI-TOF mass spectrometry and samples were dialyzed against experimental buffer (20 mM sodium phosphate, pH 7.4, 150 mM NaCl, 1 mM TCEP). Protein concentration was determined by absorbance at 280 nm and samples were frozen at -80°C until further use. Peptides corresponding to p53TAD^15-27^ were purchased from GeneCust (France) in N-terminal acetylated form and dissolved in experimental buffer. Concentration of the peptides was determined by measuring A280.

### Fluorescence polarization experiments

Affinity between p53TAD and its corresponding MDM2 was determined in a fluorescence polarization assay. First, the affinity between FITC-labeled p53TAD_Human_^15-26^ and all MDM2s were determined by keeping FITC-p53TAD_Human_^15-26^ at low concentration (15 nM) and measuring FP at increasing concentrations of MDM2. A one-site binding model was fitted to these data to obtain *K*_D_ for FITC-p53TAD_Human_^15-26^ for each MDM2 variant. Next, to determine the affinity for native and non-native interactions, a complex between MDM2 (0.2 μM for *H.sapiens*, 0.5 μM for Actinopterygii AltAll, 1 μM for *L.oculatus*, *A.anguilla*, *C.harengus*. *D.rerio*, Teleost ML and Teleost AltAll, 1.5 μM for *C.Argus*, *D.rerio* and Actinopterygii ML, and 2 μM for *M.zebra*, respectively) and FITC-p53TAD_Human_^15-26^ (15 nM) was displaced by increasing concentrations of various short p53TAD^15-27^ peptides or p53TAD^Full^ ^length^ (**Supplementary Prism File 1**). From the displacement curve and *K*_D_ for FITC-p53TAD_Human_^15- 26^ we calculated *K*_D_ values for the native and non-native interactions for “displacer” peptides as described by Nikolovska-Coleska *et al*. (42) (**Supplementary Spreadsheet 6**). Parameters from binding experiments were obtained by curve fitting in GraphPad Prism 9 (**Supplementary Prism File 1**).

### ColabFold predictions

We used ColabFold (22) to predict the interactions between p53TAD^15-27^ peptides or p53TAD^Full^ ^length^ and the respective native MDM2. Only the highest confidence model was analyzed (**Fig. S9**)

## Acknowledgments

This work was funded by the Swedish Research Council (2020–04395) and the Knut and Alice Wallenberg foundation (2015.0069) to PJ, by the Swedish Research Council (2017–02907) and the Knut and Alice Wallenberg Foundation (KAW 2016.0361) to LA, and by the National Science Foundation (NSF) to RBR (DEB-1932759 and DEB-2225130) and DA (DEB-2015404 and DEB-2144325).

## Author contributions

Conceptualization and writing original draft: FM and PJ. Experiment and analysis: FM, DA, MEP, PF, EA, LA, RB and PJ.

## Supplementary information

**Figure S1.**
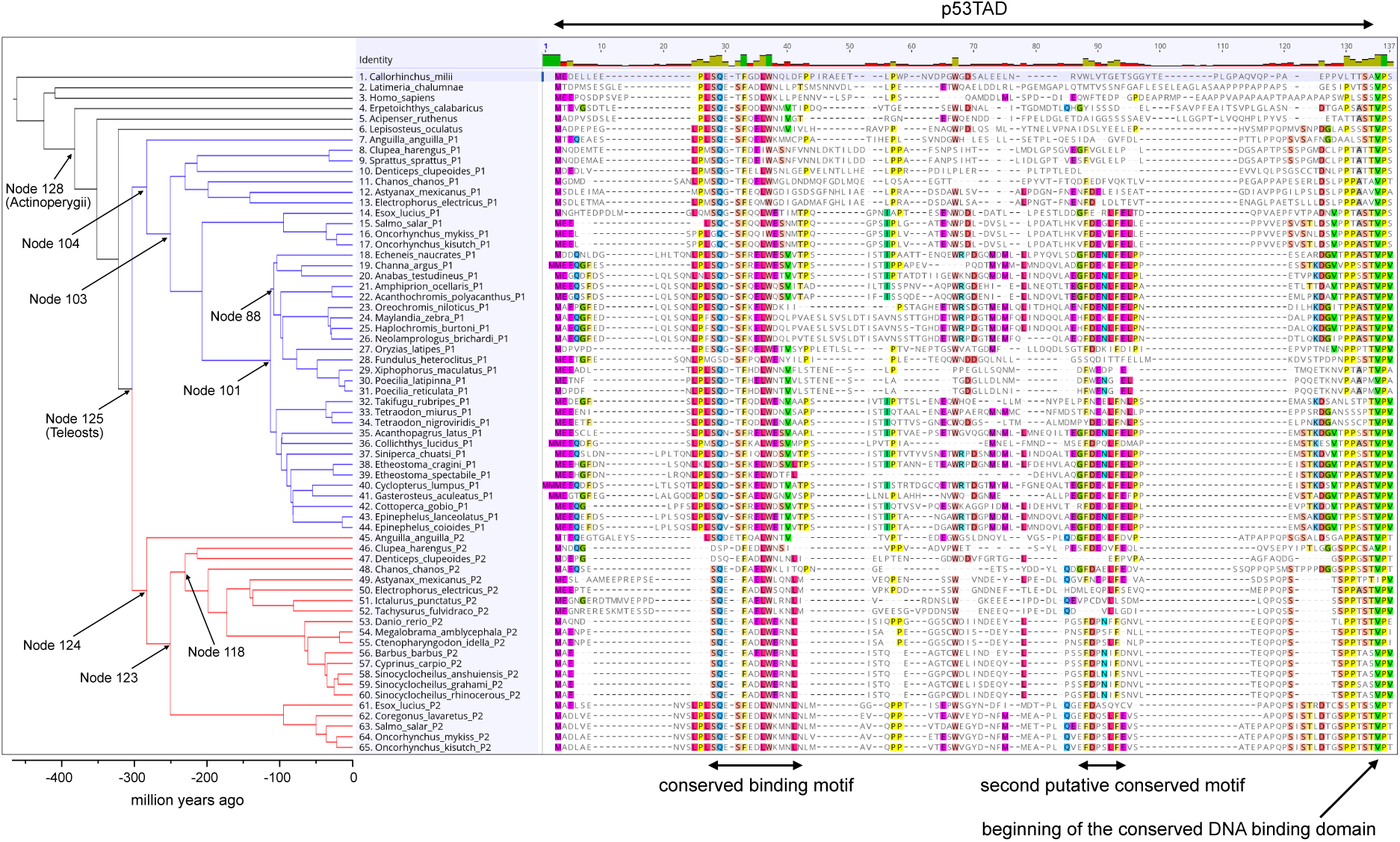
Sequence alignment of the TAD domain from p53. The sequence alignment was used for reconstruction of ancestral fish p53TAD binding motifs.

**Figure S2.**
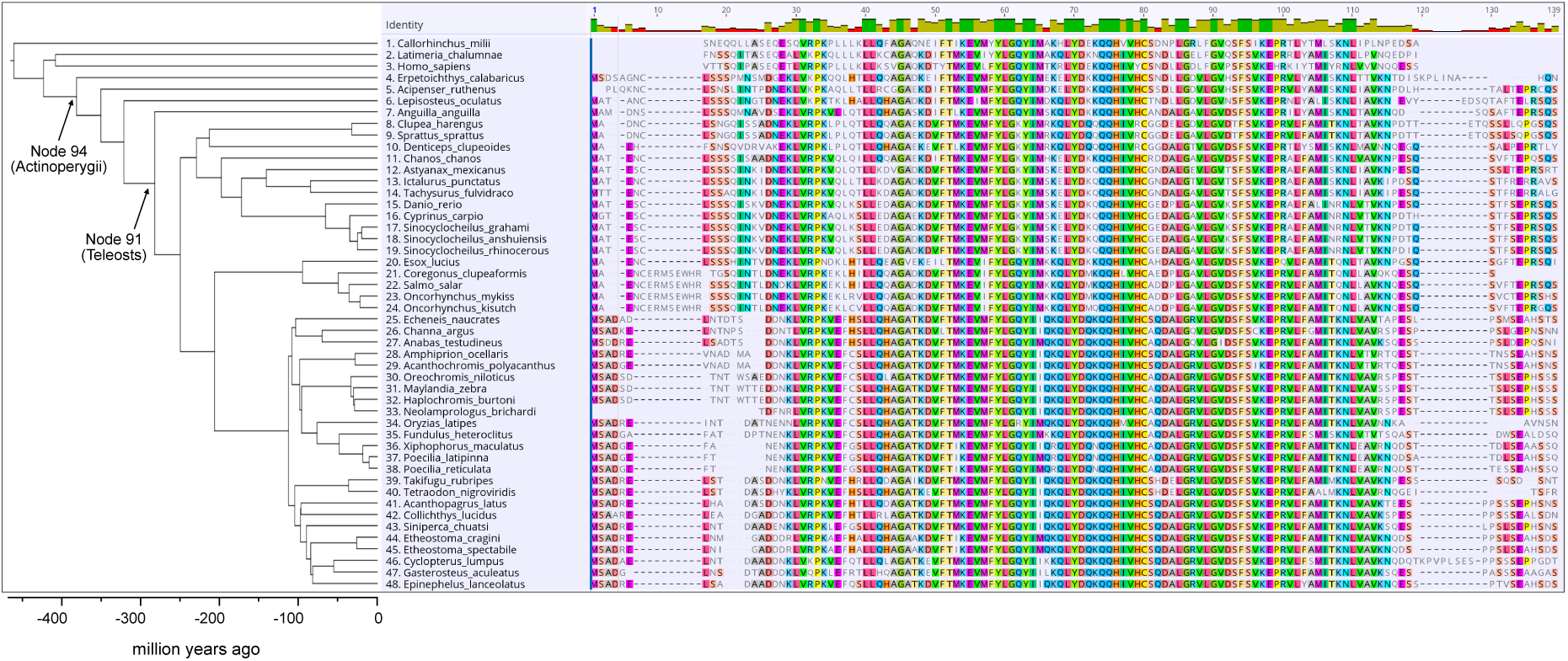
Sequence alignment of the SWIB domain from MDM2. The sequence alignment was used for reconstruction of ancestral fish MDM2 SWIB domains.

**Figure S3.**
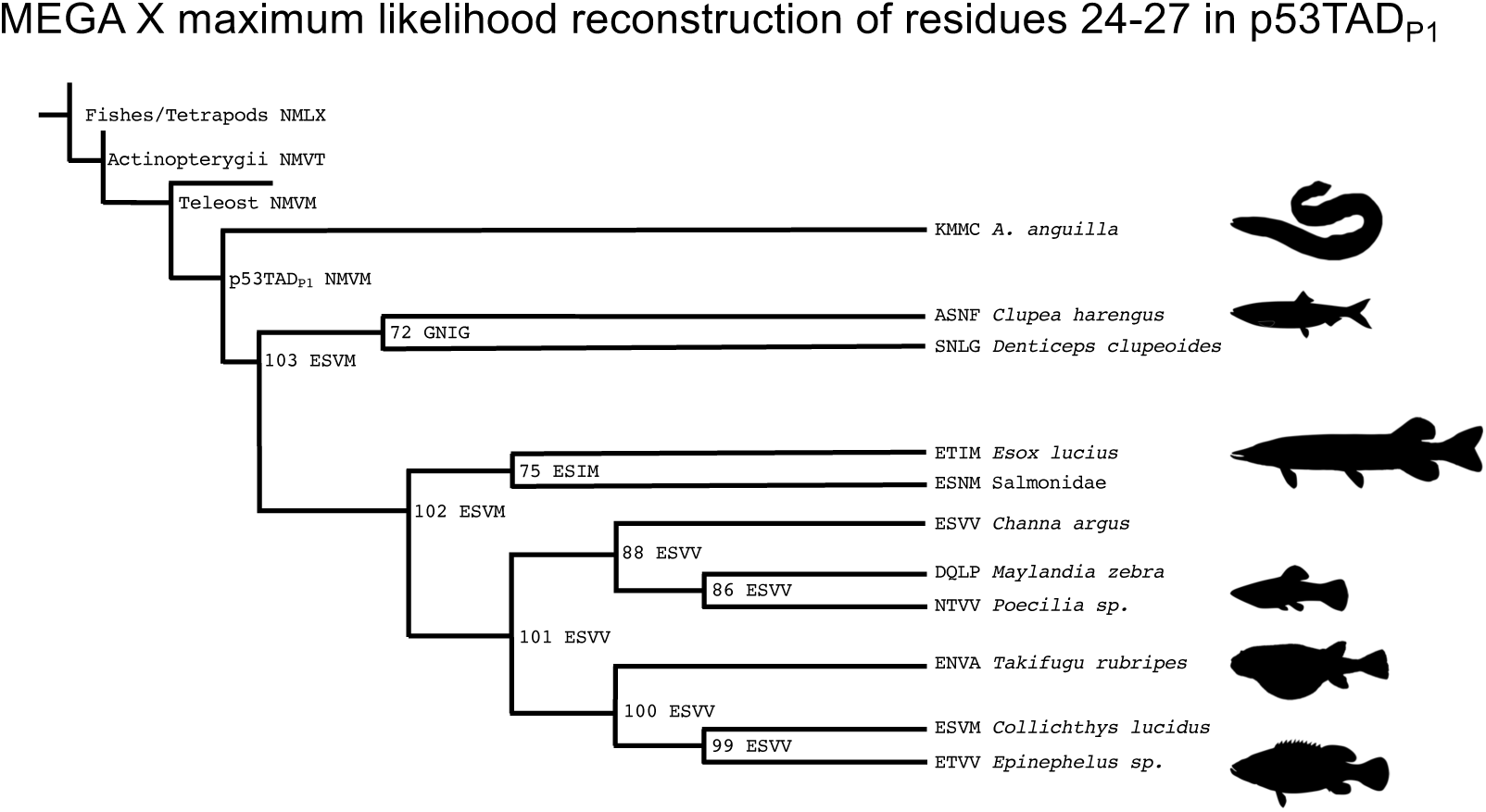
Evolution of p53TAD ^24-27^. The maximum likelihood residues for residues 24-27 for nodes in the phylogenetic tree in Fig. 2 together with selected extant sequences. See Supplementary Spreadsheet 2 and 3 and Supplementary text file 1 for posterior probabilities and AltAll residues. Pictures are from Phylopic.org.

**Figure S4.**
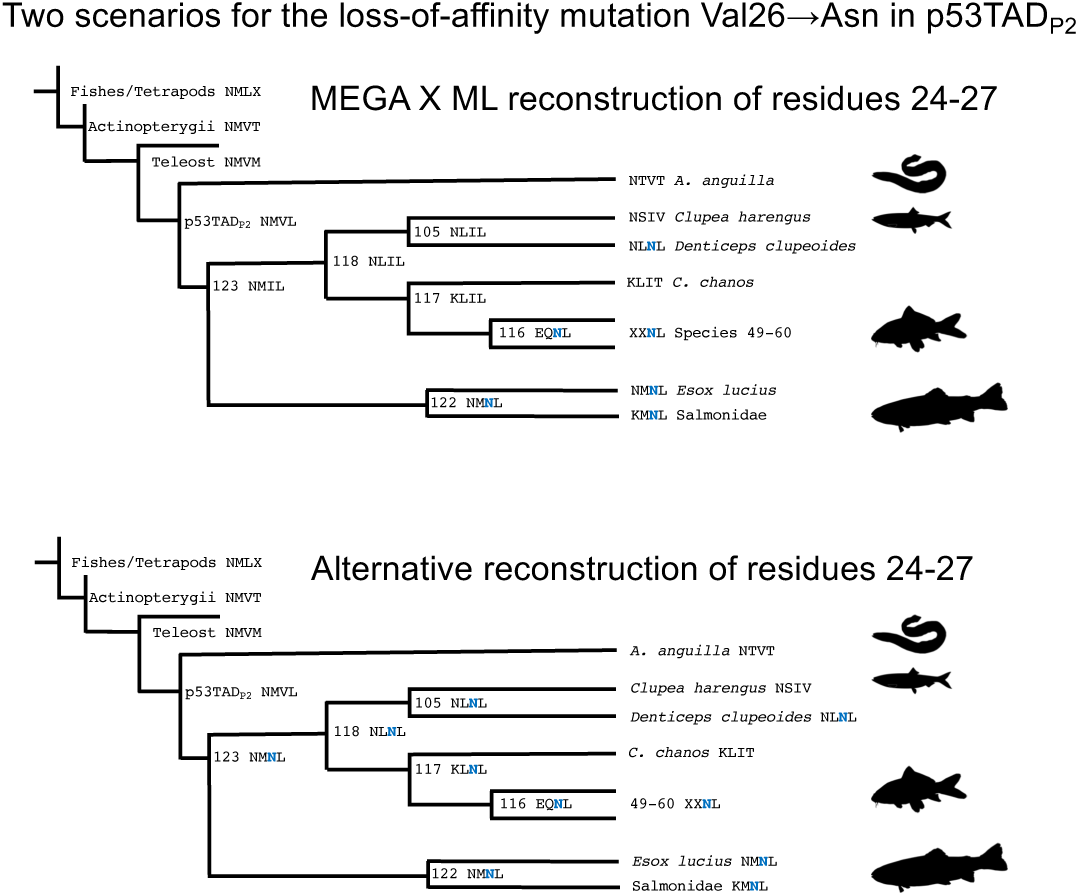
Evolution of p53TAD_P2_^24-27^. The maximum likelihood residues for residues 24-27 for nodes in the phylogenetic tree in Fig. 2 together with selected extant sequences. See Supplementary Spreadsheet 2 and 3 and Supplementary text file 1 for posterior probabilities and AltAll residues. Two different scenarios are shown, (*i*) the one reconstructed by MEGAX where the first mutation is Val_26_→Ile, and where Asn (blue N) is introduced later at three different occasions, and (*ii*) an alternative scenario where Val_26_→Asn occurs already at node 103 and Ile_26_ is introduced later in separate lineages. In both cases convergence is observed, suggesting that certain (combinations of) residues are preferred for optimal fitness. Pictures are from Phylopic.org.

**Figure S5.** Sequence alignment by affinity. The p53TAD peptides with highest affinity are at the top and those with lowest at the bottom. The order is based on (A) affinity for the native MDM2 and (B) affinity for human MDM2.

**Figure S6.**
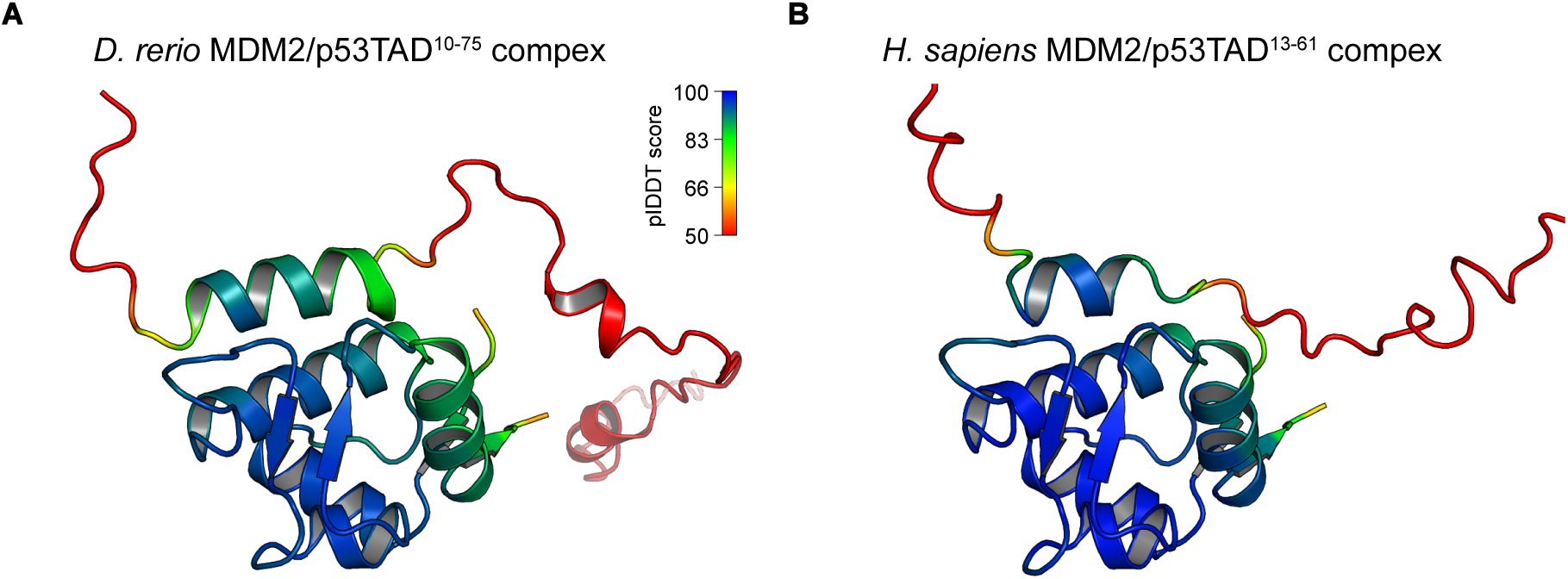
Colabfold prediction of the MDM2/p53TAD^Full^ ^length^ complexes. **A)** prediction for *D. rerio* complex. Models are colored in spectra in accordance with the prediction confidence score (pIDDT) with blue signifying high prediction confidence and red low confidence (pIDDT<50). **B)** prediction for *H. sapiens* complex. Coloring as in B). Note a significantly shorter helix of the bound p53 binding motif for the human complex in comparison to *D. rerio* complex.

**Figure S7.**
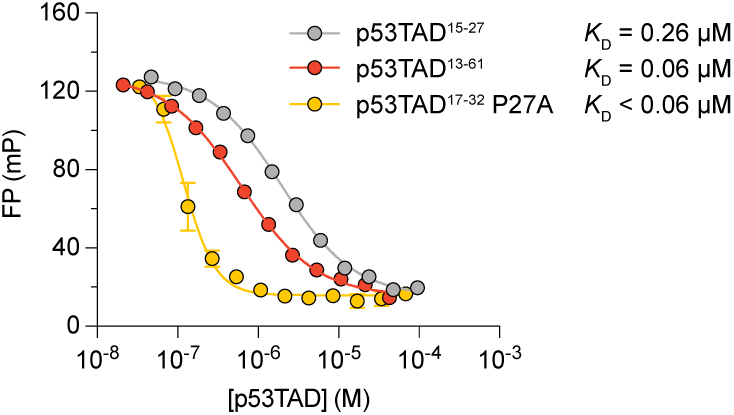
High affinity of human p53TAD^17-32^ P27A mutant. Fluorescence polarization experiment showing displacement of the labeled probe by human p53TAD^15-27^ (short), p53TAD^13-61^ (full length) and p53TAD^17-32^ P27A mutant. Note that the *K*D of the p53TAD^17-32^ P27A mutant is too low to be determined in our fluorescence polarization assay, likely in the low to sub nM range.

**Figure S8.**
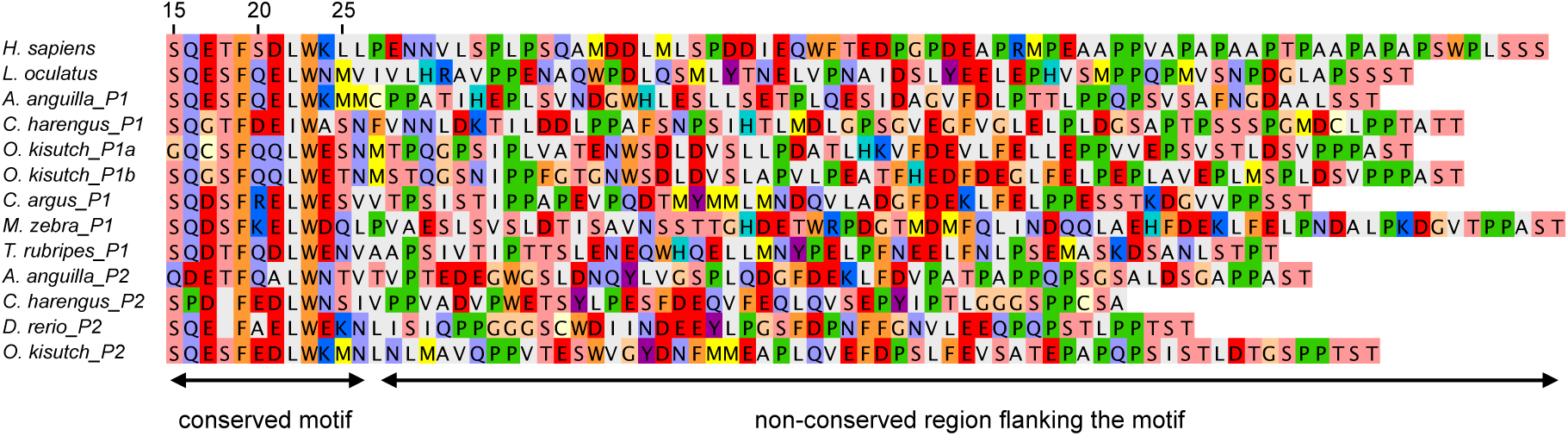
Non-conserved nature of the C-terminal flanking region. The transactivation domains of extant fish p53TAD constructs used in this study were aligned at the conserved binding motif (without the N-terminal part). The C-terminal flanking regions are “aligned” without gaps to showcase the variability and lack of clear patterns in the sequence following the conserved Trp_23_. Note that the motif with the highest affinity (*A. anguilla* P1) has methionine both at position 25 and 26.

**Figure S9.**
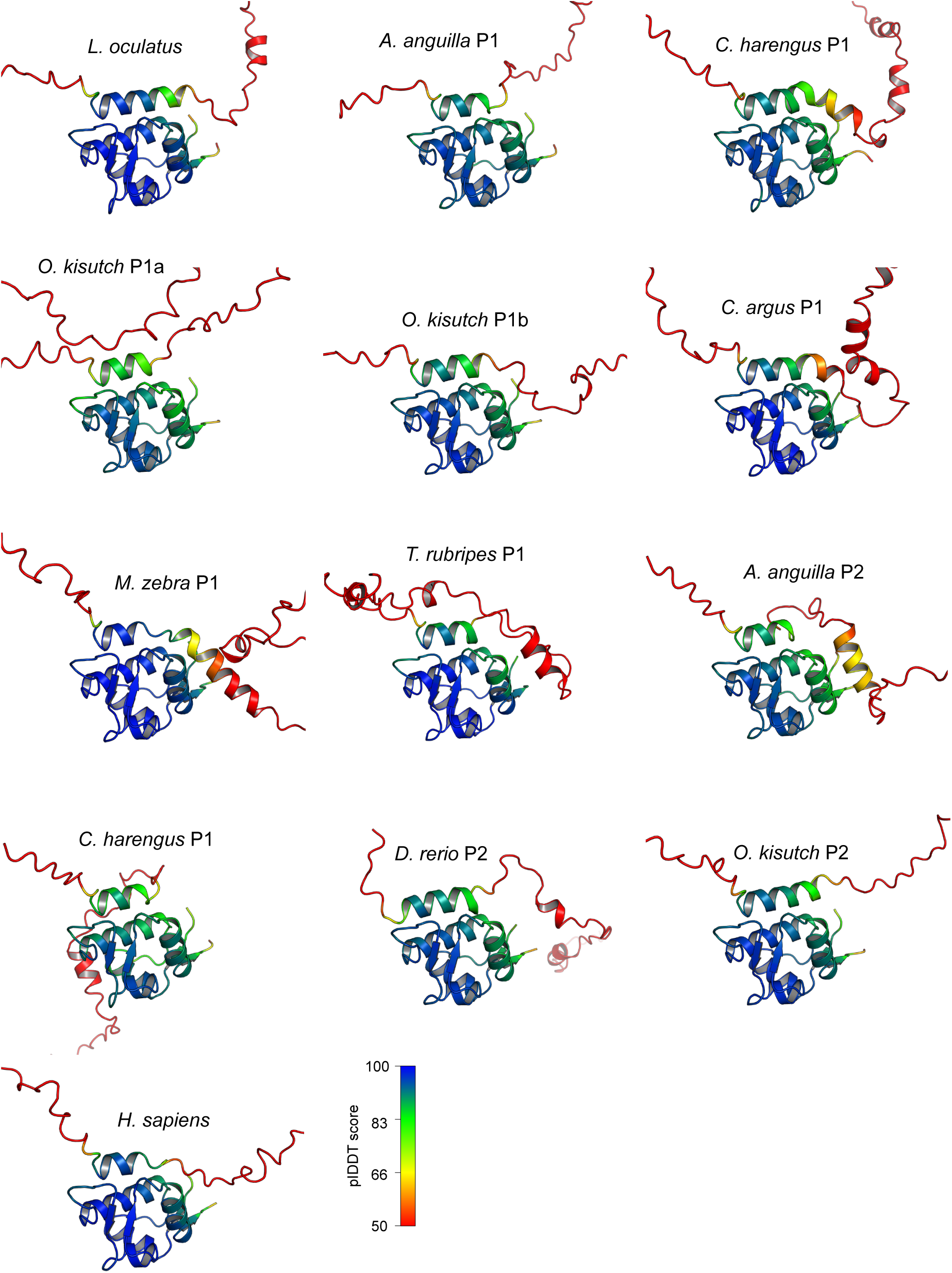
Colabfold prediction for all present day MDM2/p53TAD^full^ ^length^ interactions investigated in this study. The structures are colored according to the confidence score of the prediction, blue is highest confidence and red lowest.

**Supplementary Spreadsheet 1.** Ancestral sequence reconstruction of nodes in the p53TAD tree (**Fig. 2**).

**Supplementary Spreadsheet 2.** Ancestral sequence reconstruction of nodes in the MDM2 SWIB tree (**Fig. S1**).

**Supplementary Spreadsheet 3.** Reconstructed maximum likelihood (ML) and AltAll sequences for p53TAD used in the study. The number in bold refers to the node in the tree and other numbers are posterior probabilities from the reconstruction (**Supplementary Spreadsheet 1**).

**Supplementary Spreadsheet 4.** Reconstructed maximum likelihood (ML) and AltAll sequences for MDM2 SWIB domain used in the study. Residue numbers, node numbers and posterior probabilities from the reconstruction (**Supplementary Spreadsheet 2**) are shown.

**Supplementary Spreadsheet 5.** Sequences of p53TAD peptides and MDM2 SWIB domains used in the study.

**Supplementary Spreadsheet 6.** Calculated *K*_D_ from fluorescence polarization displacement experiments.

**Supplementary Prism File 1.** All fluorescence polarization experiments performed in this study. Concentrations of the “displacer” peptides can be found for each displacement experiment.

**Supplementary Table 1.**
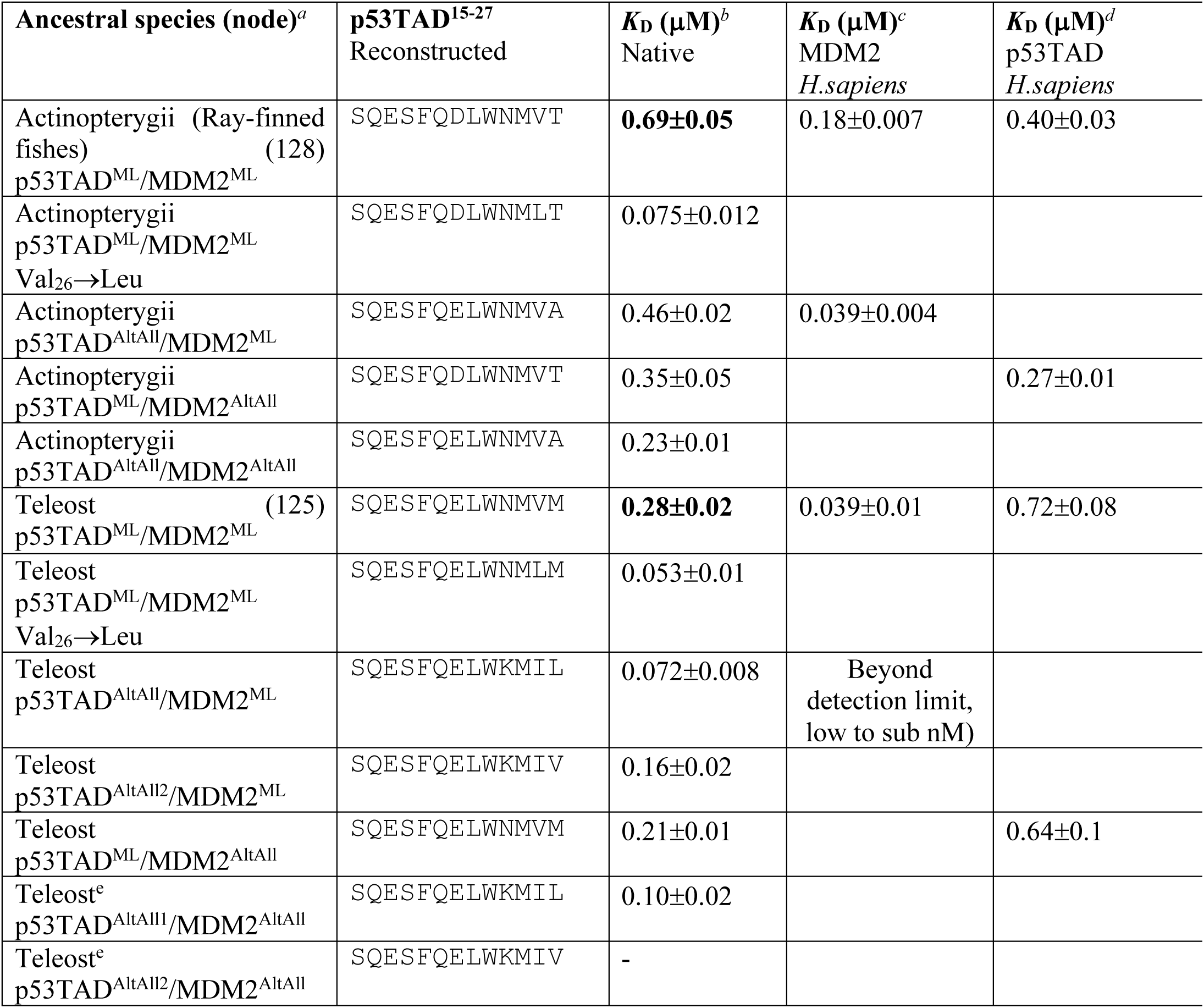
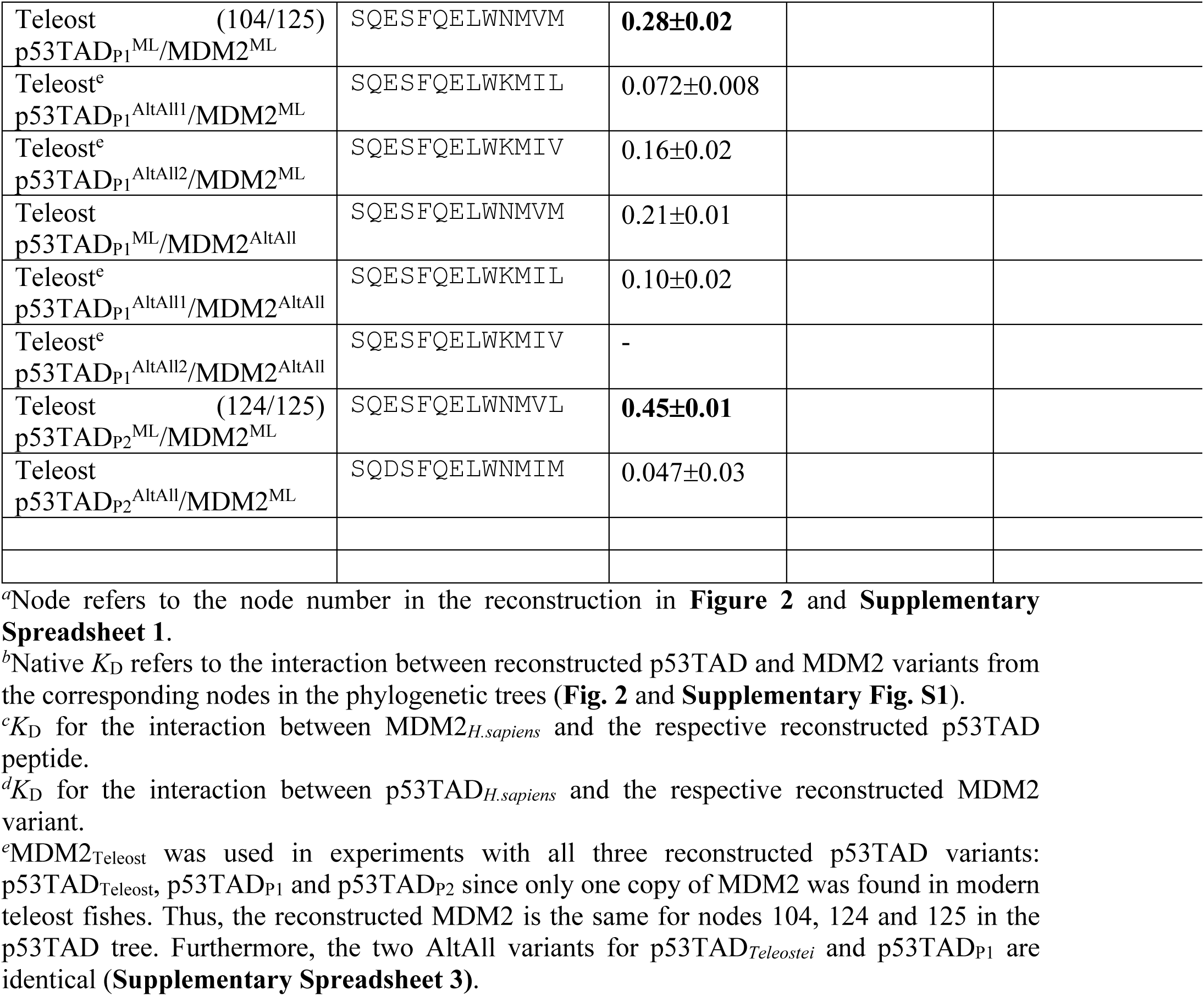
Affinities between ancestral p53TAD^15-27^ and MDM2. The complex between labeled p53TAD_Human_^15-27^ and reconstructed MDM2_Actinopterygii_ or MDM2_Teleost_ was displaced by unlabeled reconstructed p53TAD^15-27^ from the different evolutionary nodes. Native maximum likelihood *K*_D_ values are in bold.

